# Motor neuron activity induces proteomic stress in the target muscle

**DOI:** 10.1101/2023.08.31.555699

**Authors:** Saurabh Srivastav, Kevin van der Graaf, Prisha C. Jonnalagadda, Maanvi Thawani, James A. McNew, Michael Stern

## Abstract

Several lines of evidence demonstrate that increased neuronal excitability can enhance proteomic stress. For example, epilepsy can enhance the proteomic stress caused by the expression of certain aggregation-prone proteins implicated in neurodegeneration. However, unanswered questions remain concerning the mechanisms by which increased neuronal excitability accomplishes this enhancement. Here we test whether increasing neuronal excitability at a particular identified glutamatergic synapse, the Drosophila larval neuromuscular junction, can enhance the proteomic stress caused by mutations in the ER fusion gene *atlastin* (*atl*). It was previously shown that larval muscle from the *atl^2^* null mutant is defective in autophagy and accumulates protein aggregates containing ubiquitin (poly-UB aggregates). To determine if increased neuronal excitability might enhance the increased proteomic stress caused by *atl^2^*, we activated the *TrpA1*-encoded excitability channel within neurons. We found that TrpA1 activation had no effect on poly-UB aggregate accumulation in wildtype muscle, but significantly increased poly-UB aggregate number in *atl^2^*muscle. Previous work has shown that *atl* loss from either neuron or muscle increases muscle poly-UB aggregate number. We found that neuronal TrpA1 activation enhanced poly-UB aggregate number when *atl* was removed from muscle, but not from neuron. Neuronal TrpA1 activation enhanced other phenotypes conferred by muscle *atl* loss, such as decreased pupal size and decreased viability. Taken together, these results indicate that the proteomic stress caused by muscle *atl* loss is enhanced by increasing neuronal excitability.

## Introduction

Autophagy defects leading to accumulation of protein aggregates, often containing ubiquitin, are hallmarks of several prominent neurodegenerative disorders such as Alzheimer’s, Parkinson’s, and Huntington’s disease [1–3]. For example, mice defective in autophagy exhibit neurodegeneration [4, 5], whereas inducing autophagy via treatment with rapamycin could partially suppress aggregate formation in model systems [6, 7]. These data suggest that a common feature of neurodegenerative disorders might be proteomic stress, caused by any number of cell stressors including expression of aggregation-prone proteins or inhibition of autophagy.

Proteomic stress likewise appears to be involved in the family of neurodegenerative disorders termed the “Hereditary Spastic Paraplegias” (HSPs), which are caused by mutations in any of at least 72 genes. These HSP mutations confer the common pathology of progressive spasticity and paralysis, likely due to degeneration of upper motor neurons. Analysis of several HSP genes implicate defective autophagy as a common causal factor. For example, the HSP genes *spastizin (SPG15), strumpellin (SPG8)* and *Spatacsin (SPG11)* encode proteins critical for autophagy function [8–10]. In addition, the HSP gene *atlastin (SPG3A)* encodes an ER fusion protein that binds to the autophagy inducer ULK1 to induce autophagosome formation [11]. Drosophila muscle lacking *atl* accumulates undegraded autophagy cargo as well as protein aggregates containing ubiquitin (poly-UB aggregates) and exhibits precocious degeneration [12, 13]. Taken together, these data raise the possibility that defective autophagy is a causal factor for protein aggregate accumulation and ultimately degeneration.

Several lines of evidence demonstrate a link between neurodegeneration and conditions such as epilepsy that increase neuronal excitability. For example, neurodegeneration can cause seizures [14] whereas decreasing Tau expression can suppress seizures [15]. Likewise, increasing neuronal excitability can exacerbate degeneration. For instance, temporal lobe epilepsy can induce tauopathy, whereas increasing neuronal excitability can exacerbate tauopathy [16, 17]. Furthermore, hyperexcitability is observed early in Alzheimer’s progression in both patients and in mouse models [18, 19]. Increased synaptic activity and consequent glutamate toxicity could represent one mechanism by which hyperexcitability could induces degeneration [20–22]. However, the mechanisms linking neuronal excitability with proteomic stress and ultimately degeneration remain incompletely understood.

Here we examine the effects of increased neuronal excitability, accomplished by activating the heat-inducible excitability gene *TrpA1* [23] within motor neurons, on proteomic stress at the glutamatergic Drosophila larval neuromuscular junction. We found that motor neuron TrpA1 activation in otherwise wildtype larvae failed to induce poly-UB aggregate accumulation in target muscle. However, activating TrpA1 in motor neurons from *atl^2^* null larvae, as well as from larvae in which *atl* was knocked down in muscle, significantly enhanced accumulation of these poly-UB aggregates. Expressing a second excitability gene, *Sh^DN^*, likewise enhanced poly-UB aggregate accumulation in *atl^2^* larval muscle. We conclude that increasing motor neuron excitability increases proteomic stress in the target muscle, which is revealed in muscle in which autophagy machinery is crippled by *atl* loss.

## Materials and Methods

### Drosophila stocks and media

The following *s*tocks were obtained from the Bloomington Drosophila stock center (BDSC, Bloomington, Indiana): *4EBP-lacZ* (#9558), *UAS-Dcr-2* (#24650), *24B-Gal4* (#1767), *Df(atl)/TbTM6* (#7948) *OK371-Gal4* (#26160) and *nSyb-LexA (attP2,* #51951*), UAS-TrpA1* (*attP16*, #26263). Flies bearing *UAS-Sh^DN^* were generously provided by Tim Mosca (Thomas Jefferson University), flies bearing *Aop-TrpA1* in the *VK05* site were generously provided by Gerry Rubin (Janelia HHMI) and Kristin Scott (University of California, Berkeley). *UAS-atlRNAi* and *atl^2^* were described previously [24]. All fly stocks were maintained on standard cornmeal/agar media as previously described [25]. All larvae were collected for imaging at the wandering third instar stage. For experiments shown in Figures 1-4 and in all supplemental figures, larvae were reared at 28 ° C for all stages. For experiments shown in Figures 5 and 6, eggs were collected and reared for two days at room temperature, then transferred to 28 ° C until they were collected for analysis. For experiments shown in Figure 7, larvae were reared at room temperature (21 ° C –22 ° C) for all stages.

**Figure 1.**
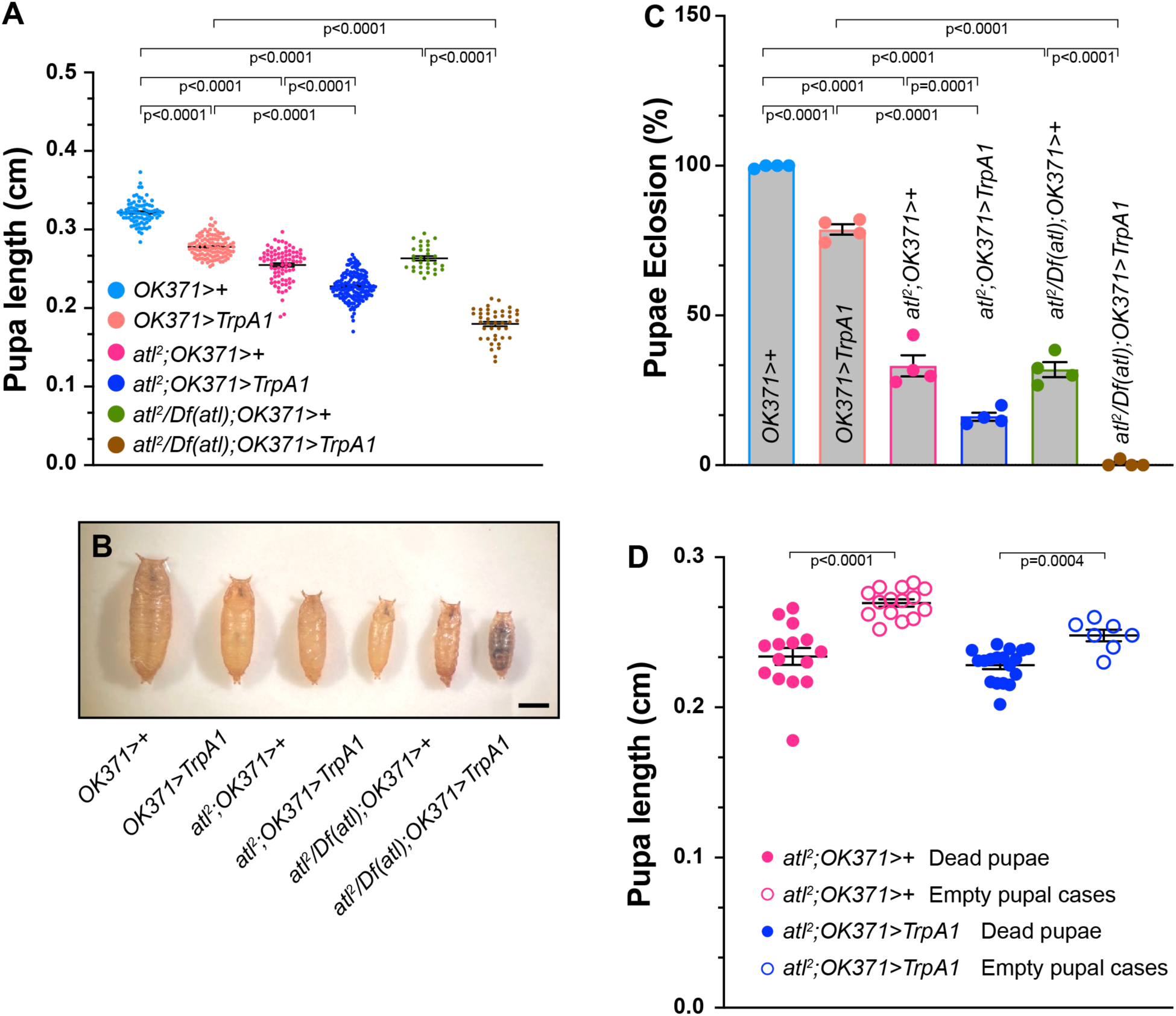
– Enhancement of *atl^2^* growth and viability phenotypes by TrpA1 activation. A) Pupal length (Y axis) from the indicated genotypes (color-coded, shown on X-axis) was measured with a dissecting stereomicroscope. Pupal length shown as a scatterplot. Means indicated by horizontal lines. B) Representative images of pupae from the indicated genotypes (color-coded). Scale bar = 0.1 cm. C) Eclosion frequency (Y axis) of pupae from the indicated genotypes (X axis). Means+/-SEMS are shown. Genotypes were color-coded as in panel A and B. D) Lengths of empty pupal cases (open circles) and pupae containing dead animals (closed circles) of the indicated genotypes, shown as a scatterplot. Means +/-SEMs are indicated. One-way ANOVA and Tukey’s post-hoc test was used for statistical analysis shown in panels A and C. Student’s t-test was used for statistical analysis shown in panel D.

### Immunocytochemistry

Larvae were dissected on Sylgard plates in 1x PBST (0.3% Tween 20 [Fisher Scientific, BP337-500] in 1x phosphate-buffered saline [Gibco, 10010-023]). Dissected larvae were fixed by treating with 4% paraformaldehyde (Electron Microscopy Sciences, 50-980-487) for 10 min at room temperature (RT). Fixed larvae were washed three times with PBST for 10 min each time, then blocked by treating the larval samples with 1% bovine serum albumin (BSA; MP-Biomedicals, 0218062080) for 30 min. Samples were incubated with primary antibodies overnight at 4℃ followed by three washes with PBST for 10 min each time. Samples were then incubated overnight at 4℃ with secondary antibody, followed by three washes with PBST. Lastly, samples were mounted on slides using Vectashield containing DAPI (Vector laboratories, H-1200-10).

Mouse anti-polyubiquitin antibody (1:1000, EMD Millipore, ST1200-100UG) was the primary antibody used in this work. The following secondary antibodies were used in this work: goat anti-mouse IgG coupled with FITC (1:1000, Abcam, ab6785).

Neuromuscular junctions were immunostained with Alexa Fluor 647 conjugated rabbit anti-horseradish peroxidase (1:1000, Jackson ImmunoResearch Laboratories, 323-605).

### Image quantitative analysis

All fluorescent images acquired were quantified and analyzed using the Surface module of Imaris v10.1. (Bitplane, Zurich, Switzerland) as previously described [13]. Poly-UB puncta number was obtained from the statistical data available for the surface created. The quantification of cytoplasmic poly-UB required an initial step in which nuclei were masked based on DAPI staining to remove inconsistent nuclear poly-UB signal.

### Construction of flies bearing *Aop-atl^+^* and *Aop-atl^K51A^*

The wildtype Drosophila *atlastin* coding sequence was generated by PCR from pJM681 [26], cut with NotI and XbaI, and ligated into pJFRC19*-13XlexAop2* (Addgene 26224) to generate pJM1181. The coding sequence for the GTPase mutant *atl^K51A^* was generated by PCR from pJM69 [26], cut with NotI and XbaI, and ligated into pJRC19 (Addgene 26224) to generate pJM1184. These constructs were introduced into embryos at the *attP2* site using with ɸC31-mediated recombination (GenetiVision, Houston TX). Insert-containing flies were recognized by eye color.

### Antibody production and Western blotting

Recombinant Drosophila atlastin (soluble domain, amino acids 1-422, pJM781 [27]) was expressed in *E. coli* and purified by nickel affinity chromatography and used as an antigen to produce rabbit polyclonal antibodies (RC77, RC78, Cocalico Biologicals).

To measure Atlastin levels by Western blotting, protein extracts were generated either from whole flies (for *atl^2^* and control) or heads (for *atlRNAi*, *atl^K51A^* and controls). Following protein extraction, proteins were quantified, size-separated by SDS-PAGE chromatography, and Western blotting was performed as described previously [13]. After transfer, membranes were blocked by treatment for 30 minutes with 5% powdered milk prepared in 1xTBST. Membranes were then incubated overnight at 4℃ with primary antibodies, followed by three washes in TBST for 10 minutes each. Membranes were then incubated with HRP-conjugated secondary antibodies for 2 hours at RT with mild shaking, followed by three washes in TBST for 10 minutes each. Membranes were then incubated with ECL reagent and observed in a LAS 4000 gel imager (Fujifilm). The following primary antibodies were used: rabbit anti-Atlastin polyclonal antibody (1:1000, RC78), and mouse anti-*Drosophila* β-tubulin antibody (1:1000, E7, DSHB). The secondary antibodies used were HRP-conjugated goat anti-mouse antibody (1:5000, #610-1319, Rockland), and HRP conjugated goat anti-Rabbit IgG (H+L) antibody (1:5000, Invitrogen, 31460).

### Pupal length measurements

Pupal images were acquired with an Olympus trinocular microscope. ImageJ (National institutes of Health) was used to measure lengths along the central axis beginning with the anterior end.

### Graphing and statistical analysis

All data analyzed in the present study were plotted and statistically analyzed using GraphPad Prism v9.3.1. One-way ANOVA with Tukey post-hoc analysis was used for all statistical analysis presented except for Figure 1D, for which Student’s t-test was used.

## Results

### Enhancement of *atl^2^* muscle phenotypes by motor neuron TrpA1 activation

Several lines of evidence suggest a link between neuronal excitability and neurodegeneration, but the mechanisms linking these processes remain incompletely understood. We addressed this question by examining effects of increasing neuronal excitability on Drosophila lacking the ER fusion protein Atlastin (Atl) [26], which accumulate undegraded autophagic cargo and ubiquitinated protein aggregates (poly-UB aggregates) in larval muscle [13], and exhibit precocious degeneration of adult muscle [12]. We increased excitability in larval motor neurons by driving expression of *TrpA1,* which encodes a heat-inducible cation channel, with the *OK371* motor neuron *Gal4* driver, and rearing larvae at the elevated temperature of 28°C. This protocol has been previously shown to increase neuronal excitability in larval motor neuron [28].

Larvae, pupae or adults lacking *atl* specifically in muscle exhibit a dwarf phenotype [13, 29, 30]. We determined whether TrpA1 activation affected body size in either a wildtype or *atl^2^* background by measuring pupal length. Consistent with previous reports, we found that body size was significantly decreased in either *atl^2^* or *atl/Df(atl)* pupae (Figure 1A, 1B). Furthermore, activating TrpA1 in motor neurons even in a wildtype background also significantly decreased pupal size. However, activating TrpA1 in an *atl^2^* or *atl^2^/Df(atl)* pupae decreased size significantly more than in *atl^2^*, *atl^2^/Df(atl)* or *TrpA1-* expressing pupae alone (Figure 1A, 1B). These results suggest that neuronal excitability enhances the *atl^2^* dwarf phenotype, which is one phenotype conferred by *atl* loss in the muscle.

We found that TrpA1 activation likewise enhanced the defect in pupal eclosion conferred by *atl* loss. Whereas virtually all wildtype pupae are able to eclose to adults (Figure 1C), only about 30% of *atl^2^*or *atl^2^/Df(atl)* pupae are able do so. TrpA1 activation in otherwise wildtype pupae also decreases eclosion frequency to ∼75% successful eclosion. However, activating TrpA1 in pupae lacking *atl* decreases successful eclosion even further – to about 15% in *atl^2^* pupae, and to <1% in *atl^2^/Df(atl)* pupae (Figure 1C).

We found that the decreased ability to eclose conferred by *atl^2^*was strongly correlated with decreased pupal size. In Figure 1D, we compared sizes of pupae in which eclosion successfully occurred (empty pupal case) with pupal sizes in which eclosion failed (dead pupae). We found that empty pupal cases were significantly larger than cases containing dead pupae in *atl^2^* regardless of TrpA1 activation status. Thus, there is a positive correlation between eclosion frequency and pupal size, which is consistent with the possibility that TrpA1 activation decreases eclosion frequency in *atl^2^* because TrpA1 activation further decreases pupal size in *atl^2^*.

In addition to decreased body size, larvae or adults lacking *atl* accumulate poly-UB aggregates within the muscle cytoplasm [12, 13]. To determine if TrpA1 activation would enhance the poly-UB phenotype conferred by *atl* loss, we examined effects of TrpA1 activation in muscle from wildtype, *atl^2^* and *atl^2^/Df(atl)* larvae. We found that TrpA1 activation in an otherwise wildtype background did not significantly increase cytoplasmic poly-UB aggregate number (Figure 2A, 2B and 2G). Cytoplasmic poly-UB aggregate number was not significantly affected in *atl^2^*, whereas *atl^2^/Df(atl)* conferred a modest (p=0.03) increase in aggregate number (Figure 2C, 2E and 2G). However, when TrpA1 activation was combined with either *atl^2^* or *atl^2^/Df(atl)*, poly-UB aggregate number was significantly increased over *atl^2^*, *atl^2^/Df(atl)* or *TrpA1-*expressing larvae alone (Figure 2D, 2F and 2G). Thus, TrpA1 enhances the larval muscle poly-UB phenotype conferred by *atl* loss.

**Figure 2.**
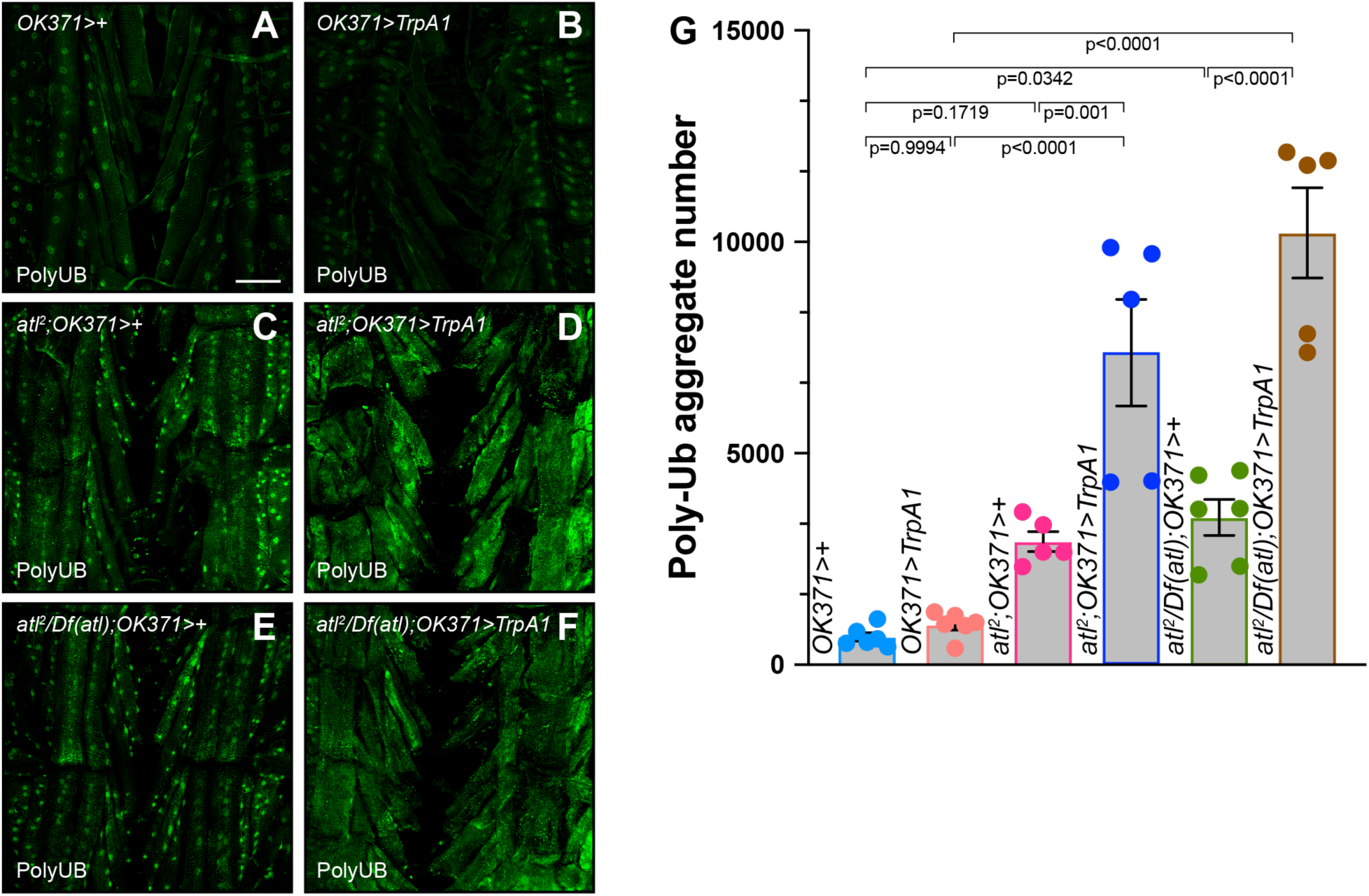
– Enhancement of *atl^2^* poly-UB aggregate phenotype by TrpA1 activation. A)-F) Confocal imaging (maximum intensity z projection) of fixed third instar larval musculature from muscle 6 stained with an anti-ubiquitin antibody (green). Scale bar = 100 µm. G) Means +/-SEMs of cytoplasmic poly-UB aggregate number (Y axis) as a function of genotype (X axis). Genotypes were color coded as shown in Figure 1A and 1B. Ubiquitin from nuclei, identified from the DAPI channel, was removed prior to quantification. One-way ANOVA and Tukey’s post-hoc test was used for statistical analysis.

### TrpA1 activation does not enhance phenotypes conferred by neuronal *atl* loss

Various muscle properties are affected by *atl* loss specifically in muscle (cell autonomous processes) whereas others are affected by *atl* loss specifically in neurons (synaptically-driven processes). For example, muscle *atl* loss increases the number of poly-UB-containing protein aggregates in both adult and larval muscle [12, 13]. *atl* loss specifically from neurons likewise increases the number of poly-UB-containing aggregates in adult muscle, but effects on larval muscle aggregate number have not been reported. Similarly, muscle *atl* loss is sufficient to decrease body size [13], but effects of neuronal *atl* loss on body size have not been reported. The enhancement by TrpA1 activation of the body size and poly-UB phenotypes conferred by *atl^2^* could reflect loss of *atl* from either neuron or muscle. To determine if *atl* loss in neurons was responsible for enhancement by TrpA1 activation, we used RNAi to knockdown *atl* specifically in motor neurons and determined if TrpA1 activation was still capable of enhancing phenotypes of *atl* loss.

First, we examined effects of neuronal *atl* knockdown and TrpA1 activation on body size and eclosion frequency, using the motor neuron *Gal4* driver *OK371* to express both the *atl RNAi* construct and *TrpA1*. We found, consistent with what we showed in Figure 1 above, that TrpA1 activation significantly decreased pupal size (Figure 3A and 3B). However, *atl* knockdown in motor neurons had no effect on pupal size (Figure 3A and 3B), and TrpA1 activation only slightly decreased size in *atl* knockdown pupae. These results indicate that *atl* loss in neurons is not sufficient to decrease body size and that TrpA1 activation does not interact with neuronal *atl* loss to control body size.

**Figure 3.**
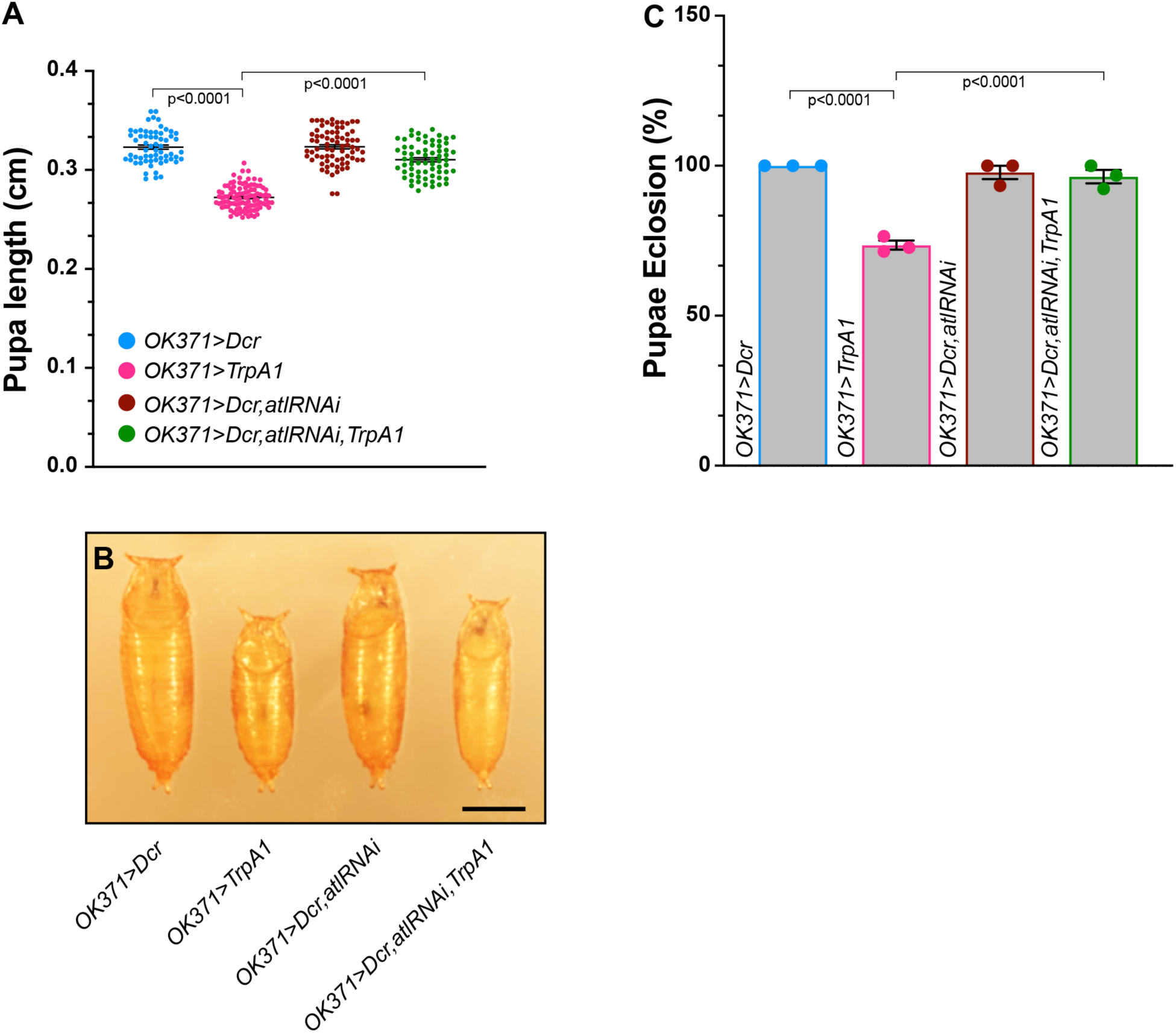
– Neuronal *atl* knockdown fails to enhance the growth and viability phenotypes of TrpA1 activation. A) Pupal length (Y axis) from the indicated genotypes (color-coded, shown on X-axis) was measured with a dissecting stereomicroscope. Pupal length shown as a scatterplot. Means indicated by horizontal lines. B) Representative images of pupae from the indicated genotypes (color-coded). Scale bar = 0.1 cm. C) Eclosion frequency (Y axis) of pupae from the indicated genotypes (X axis). Means+/-SEMS are shown. Genotypes were color-coded as in panel A and B.

Neuronal *atl* loss is likewise not sufficient to decrease pupal eclosion frequency, either in the presence or absence of TrpA1 activation (Figure 3C). Furthermore, whereas TrpA1 activation in an otherwise wildtype background decreased pupal eclosion frequency, consistent with what was shown in Figure 1, TrpA1 activation failed to decrease eclosion frequency in the context of neuronal *atl* knockdown. In fact, neuronal *atl* knockdown unexpectedly appeared to rescue the effects of TrpA1 activation on eclosion frequency, an observation that we attribute to the presence of three *UAS-*driven transgenes in these pupae, which together titrate limiting Gal4 and consequently decrease the amount of *UAS-*driven expression from these transgenes.

Second, we examined effects of neuronal *atl* loss and TrpA1 activation on muscle poly-UB aggregate accumulation. As shown in Figure 4, TrpA1 activation in an otherwise wildtype background did not significantly affect muscle poly-UB aggregate accumulation (Figure 4A, 4B and 4E), whereas neuronal *atl* knockdown increased poly-UB aggregate number significantly, albeit modestly. This result demonstrates that neuronal *atl* loss increases poly-UB aggregate number in larval as well as adult muscle, consistent with previous reports [12]. However, we found no further increase in muscle poly-UB aggregate number when we activated TrpA1 in a background of neuronal *atl* knockdown. Taken together, these results demonstrate that *TrpA1* does not enhance phenotypes conferred by neuronal *atl* loss.

**Figure 4.**
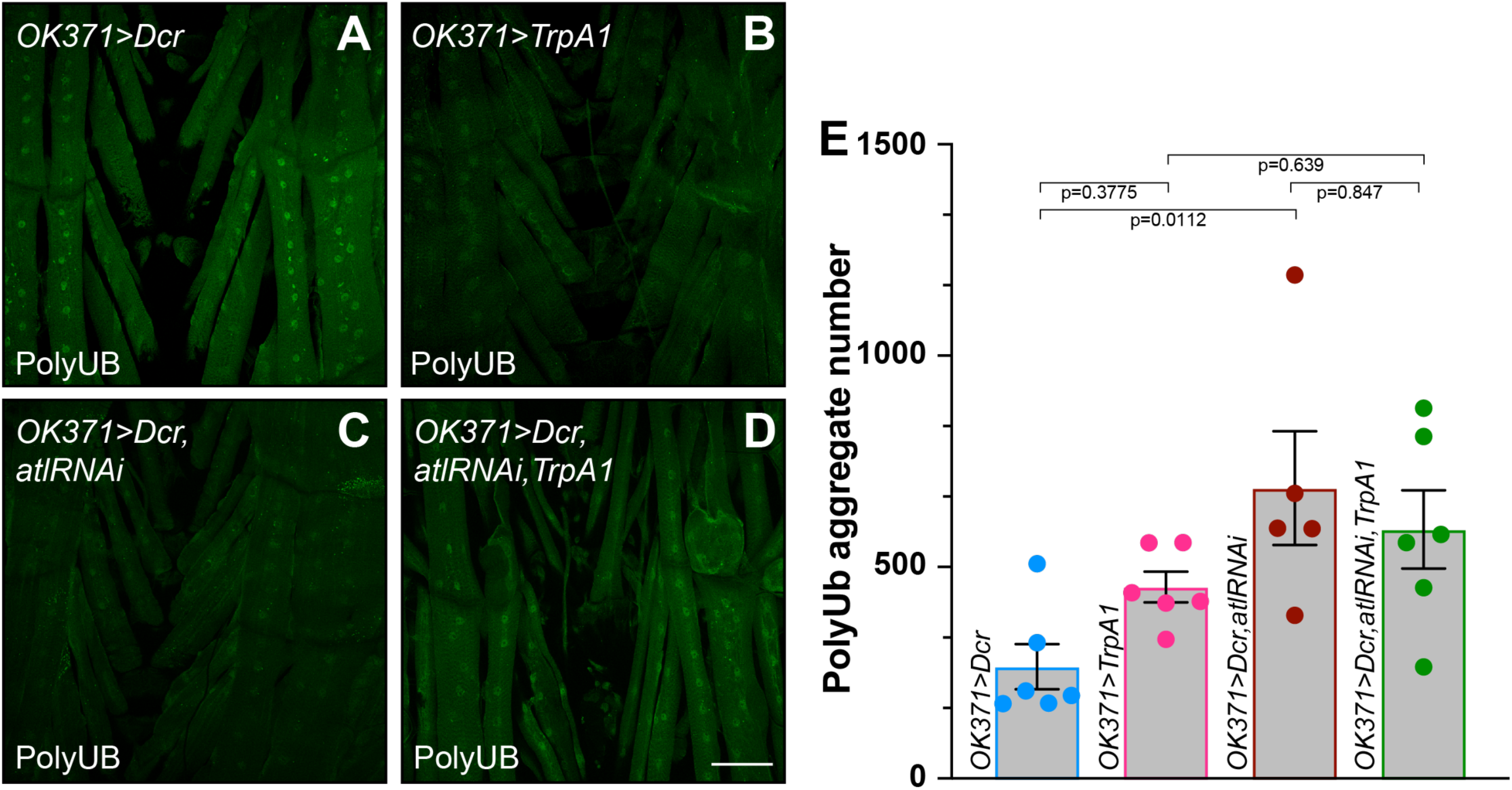
– TrpA1 activation fails to enhance poly-UB aggregate accumulation conferred by neuronal *atl* knockdown. A)-D) Confocal imaging (maximum intensity z projection) of fixed third instar larval musculature from muscle 6 stained with an anti-ubiquitin antibody (green). Scale bar = 100 µm. E) Means +/-SEMs of cytoplasmic poly-UB aggregate number (Y axis) as a function of genotype (X axis). Genotypes were color coded as shown in Figure 3A. Ubiquitin from nuclei, identified from the DAPI channel, was removed prior to quantification. One-way ANOVA and Tukey’s post-hoc test was used for statistical analysis.

### TrpA1 activation enhances phenotypes conferred by muscle *atl* loss

The observation that TrpA1 activation enhanced muscle phenotypes of *atl^2^* but not neuronal *atl* knockdown raised the possibility that this enhancement required *atl* loss from muscle. To test this possibility, we needed to independently activate TrpA1 in neurons while knocking down *atl* in muscle. To accomplish this goal, we used the *nSyb-LexA* panneuronal driver from the LexA/Aop system [31] to drive expression of *Aop-*driven *TrpA1* [32, 33]. We simultaneously used the *24B* muscle Gal4 driver to drive expression of *UAS-*driven *atl* RNAi construct, as performed previously [13].

We used two tests to validate *nSyb-LexA* as a driver capable of expressing *TrpA1* sufficiently to confer a neuronal excitability phenotype. First, we generated an antibody against Drosophila Atlastin expressed as a recombinant protein from *E. coli.* This antibody specifically recognizes Drosophila Atlastin, as the *atl^2^* deletion mutation eliminates the major band observed at 50 kd (Fig S1A). We used this antibody to compare Atlastin protein levels from heads in which either of two, *Aop*-driven constructs were expressed under the control of *nSyb-LexA*: first, an *atl* coding construct carrying the dominant-negative *atl^K51A^* allele, and second, an *atl* RNAi construct [25]. We found that *nSyb-LexA* driving *atl^K51A^* increased head Atlastin protein levels approximately 7-fold over heads from *nSyb-LexA* driving the parent empty *attP2* site (Figure S1B). Similarly, we found that *nSyb-LexA* driving *atlRNAi* decreased head Atlastin protein levels 4-fold over heads from *nSyb-LexA* driving the parent empty *attP2* site (Figure S1C). These results validate *nSyb-LexA* as an effective driver of *Aop*-driven transgenes in neurons. Note that *nSyb-LexA* driving *Aop-atl^+^* was lethal at the embryonic stage, consistent with the previous report that *elav-Gal4*-driven *atl^+^* is embryonic [24]. Second, we examined behavior of flies bearing both *nSyb-LexA* and *Aop-TrpA1* and found that etherized adults under fiber optic illumination displayed vigorous leg shaking, a hallmark of adults with increased neuronal excitability [34]. This leg shaking was similar to the leg shaking displayed by flies bearing both *OK371* and *UAS-TrpA1* but was absent from *nSyb-LexA* adults that did not carry *Aop-TrpA1.* Taken together, these observations confirm that *nSyb-lexA* driver effectively drives expression of *Aop-*driven transgenes within neurons.

We found that activating TrpA1 with the LexA/Aop system significantly decreased pupal size to an extent similar to the decrease found when TrpA1 was activated with the Gal4 system (Figure 5A and 5B). Furthermore, as described previously [13], we found that knocking down *atl* specifically in muscle also decreased pupal size. However, in contrast to the lack of enhancement when *atl* was knocked down in motor neurons, *atl* knockdown in muscle significantly enhanced the effects of TrpA1 activation on pupal size (Figure 5A and 5B). We conclude that the enhancement of the *atl* body size phenotype by TrpA1 activation requires *atl* loss in muscle but not neurons.

**Figure 5.**
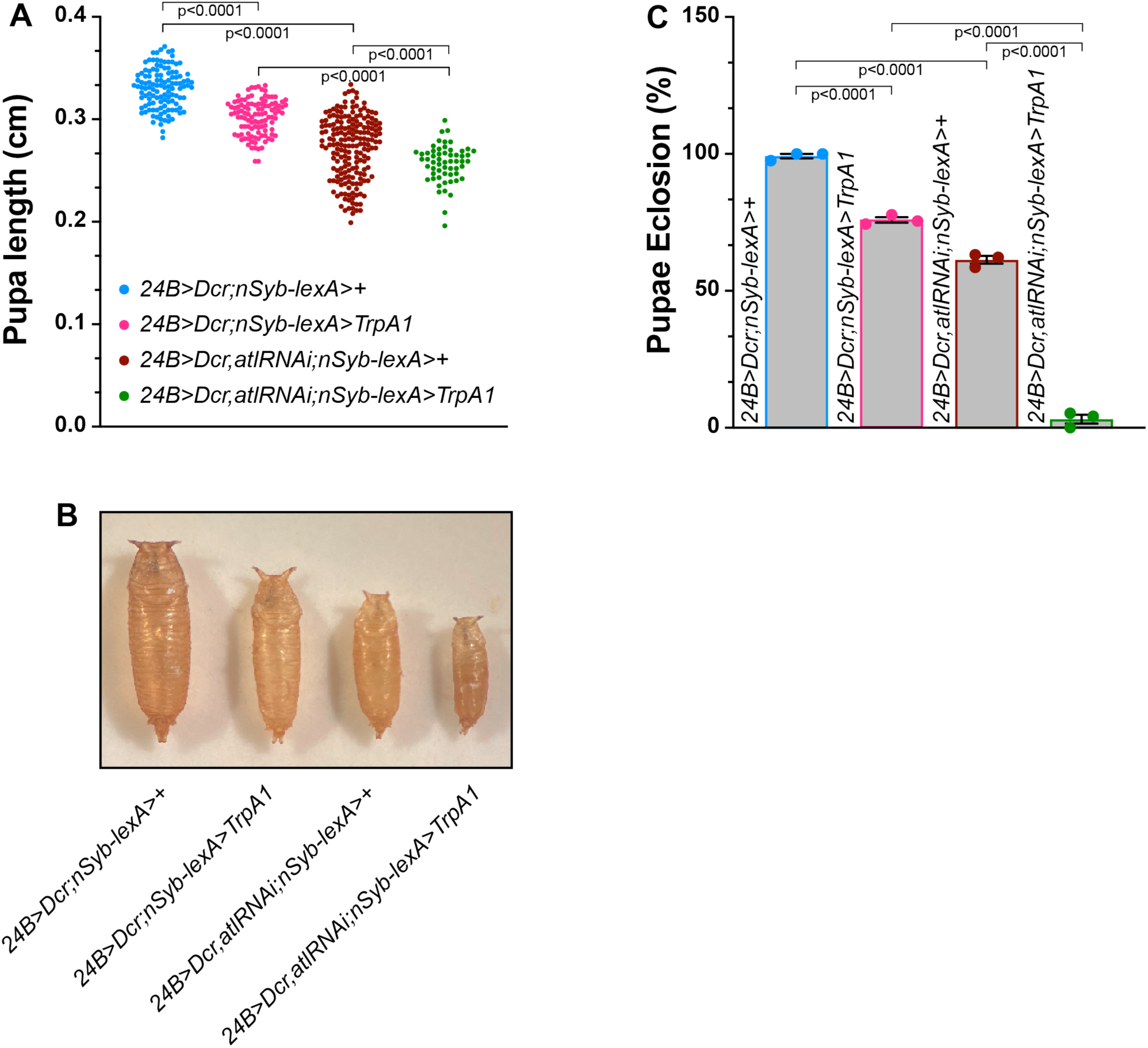
– TrpA1 activation enhances the growth and viability deficits of muscle *atl* knockdown. A) Pupal length (Y axis) from the indicated genotypes (color-coded, shown on X-axis) was measured with a dissecting stereomicroscope. Pupal length shown as a scatterplot. Means indicated by horizontal lines. B) Representative images of pupae from the indicated genotypes (color-coded). Scale bar = 0.1 cm. C) Eclosion frequency (Y axis) of pupae from the indicated genotypes (X axis). Means+/-SEMS are shown. Genotypes were color-coded as in panel A.

We also examined the effect of TrpA1 activation on eclosion frequency in muscle *atl* knockdown pupae. We found that control pupae exhibited an almost 100% frequency of eclosion (Figure 5C), consistent with what we observed for the control pupae described above. Furthermore, activating TrpA1 in an otherwise wildtype background significantly decreased pupal eclosion frequency to about 75% (Figure 5C), an observation also consistent with what we found for *Gal4-*driven *TrpA1* expression described above. Similarly, muscle *atl* knockdown in the absence of *TrpA1* expression significantly decreased eclosion frequency to about 60% (Figure 5B). However, activating TrpA1 in pupae with muscle lacking *atl* decreased successful eclosion to only about 2%. We conclude that the enhancement of the *atl* eclosion phenotype by TrpA1 activation requires *atl* loss in muscle but not neurons.

As described above, TrpA1 activation enhanced the increase in muscle poly-UB aggregate number conferred by *atl^2^* but not neuronal *atl* loss. To determine if TrpA1 activation would enhance the poly-UB phenotype conferred by muscle *atl* loss, we compared effects of TrpA1 activation on cytoplasmic poly-UB from wildtype and *atl* knockdown muscle (Figure 6). We found that TrpA1 activation in an otherwise wildtype background did not significantly increase cytoplasmic poly-UB aggregate number (Figure 6A and 6E), consistent with observations described above. Muscle *atl* knockdown in the absence of TrpA1 activity significantly increased cytoplasmic poly-UB aggregate number, consistent with previous reports [13]. When we combined TrpA1 activation with muscle *atl* knockdown, poly-UB aggregate number was significantly increased compared to either muscle *atl* knockdown or TrpA1 activation (Figure 6D and 6E). Thus, TrpA1 activation significantly enhances the larval muscle poly-UB phenotype conferred by muscle, but not neuronal, *atl* loss.

**Figure 6.**
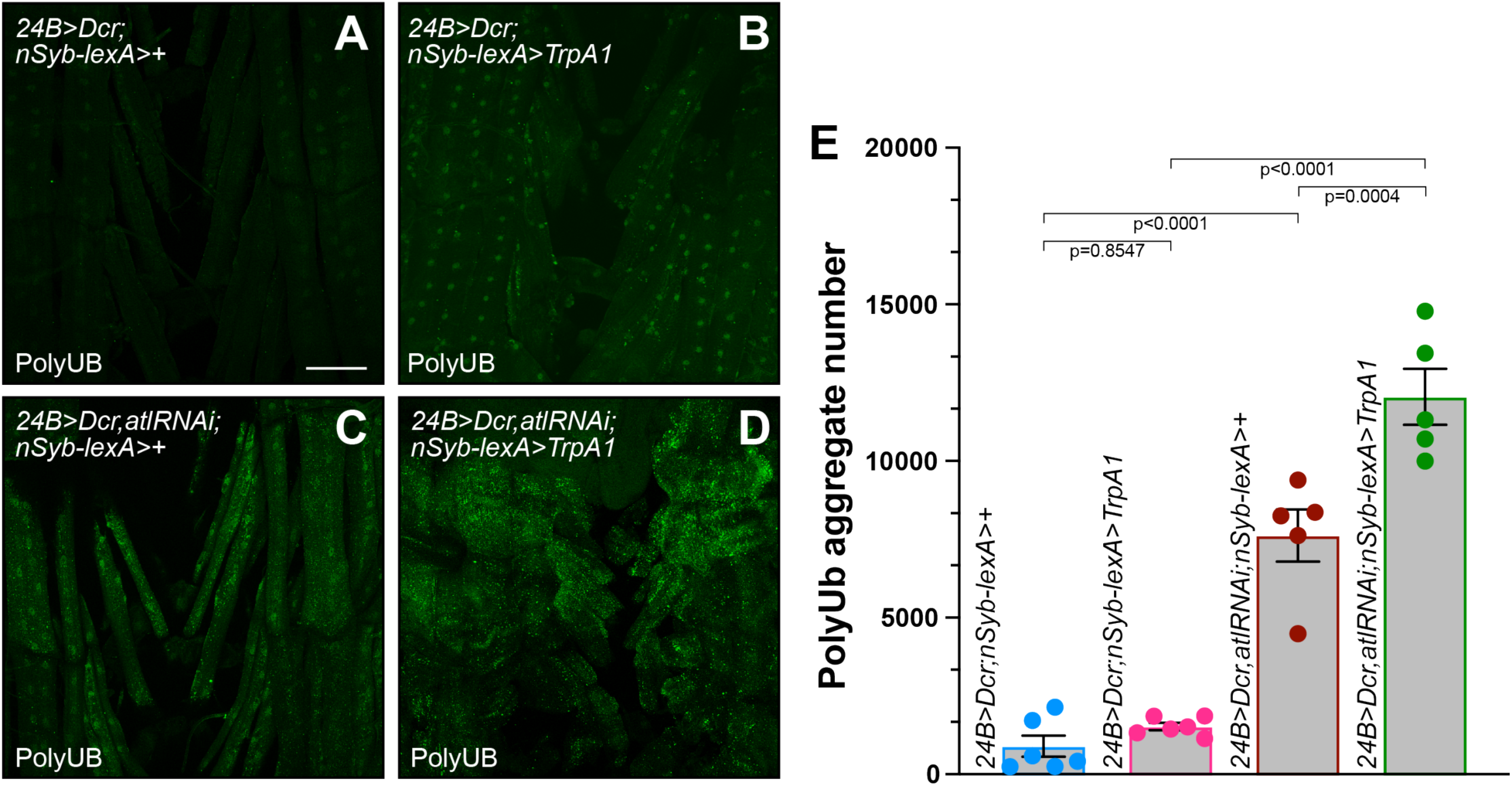
– TrpA1 activation enhances the effects of muscle *atl* knockdown on poly-UB aggregate accumulation. A)-D) Confocal imaging (maximum intensity z projection) of fixed third instar larval musculature from muscle 6 stained with an anti-ubiquitin antibody (green). Scale bar = 100 µm. E) Means +/-SEMs of cytoplasmic poly-UB aggregate number (Y axis) as a function of genotype (X axis). Genotypes were color coded as shown in Figure 5A. Ubiquitin from nuclei, identified from the DAPI channel, was removed prior to quantification. One-way ANOVA and Tukey’s post-hoc test was used for statistical analysis.

### TrpA1 activation fails to enhance *atl* effects on axon branching and synaptic bouton number

Loss of *atl* from either neurons or muscle increases axon branch and synaptic bouton number at the larval neuromuscular junction (NMJ) [13, 24]. We tested if these NMJ phenotypes could be altered by TrpA1 activation. We found that neuronal *atl* knockdown significantly increased both axon branch and synaptic bouton number (Figure S2), consistent with previous report [24]. However, TrpA1 activation had no effect on these NMJ phenotypes in a neuronal *atl* knockdown background (Figure S2). Likewise, the increased synaptic branch and bouton number conferred by muscle *atl* knockdown was not further enhanced by TrpA1 activation (Figure S2). We conclude that only certain phenotypes of *atl* loss are enhanced by increased neuronal excitability.

### Enhancement of *atl^2^* poly-UB phenotypes by a second excitability transgene

To determine if the interactions between TrpA1 activation and muscle *atl* loss on poly-UB aggregate accumulation was specific to TrpA1, or could be observed with other excitability transgenes, we tested if expressing a second excitability transgene would likewise enhance the cytoplasmic poly-UB phenotype of *atl^2^*. We chose a transgene carrying a dominant-negative mutation in *Shaker (Sh)*, which encodes a potassium channel subunit. We chose this transgene because its effects on excitability are particularly well documented [35–38]. We found that *OK371-*driven *Sh^DN^* expression in an otherwise wildtype background had no effect on larval muscle poly-UB aggregate number (Figure 7B and 7E). In contrast, this expression of *Sh^DN^* significantly enhanced the effects of *atl^2^* on aggregate number (Figure 7D and 7E).

**Figure 7.**
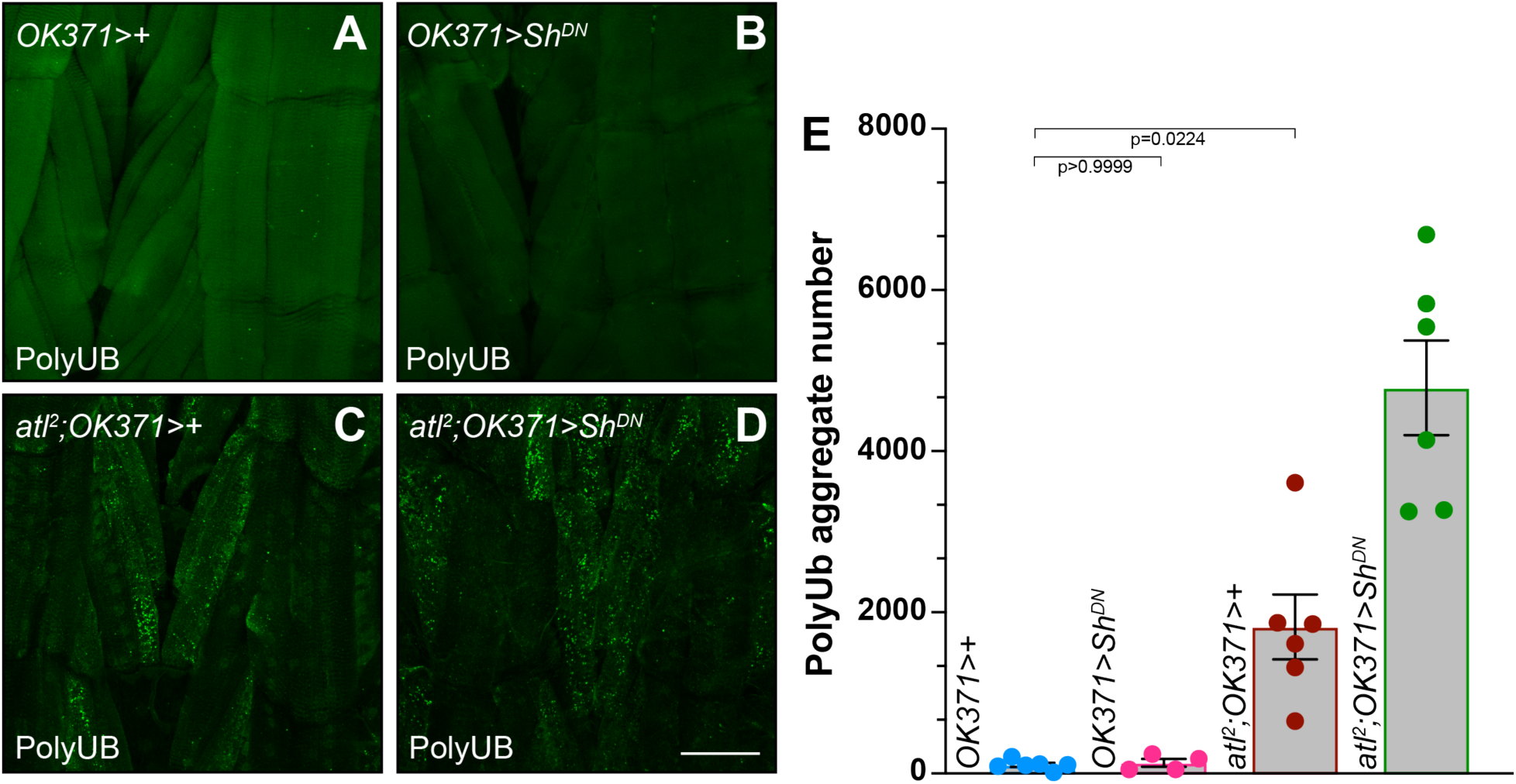
– Enhancement of the *atl^2^* poly-UB aggregate accumulation phenotype by *Sh^DN^* expression. A)-D) Confocal imaging (maximum intensity z projection) of fixed third instar larval musculature from muscle 6 stained with an anti-ubiquitin antibody (green). Scale bar = 100 µm. E) Means +/-SEMs of cytoplasmic poly-UB aggregate number (Y axis) as a function of genotype (X axis). Ubiquitin from nuclei, identified from the DAPI channel, was removed prior to quantification. One-way ANOVA and Tukey’s post-hoc test was used for statistical analysis.

## Discussion

Several lines of evidence demonstrate a link between conditions that increase neuronal excitability and proteomic stress [16, 39, 40]. However, neither the causal relationship between these phenomena nor the mechanisms underlying this linkage are completely understood. Here we use the glutamatergic Drosophila larval neuromuscular junction to address these questions. We increased neuronal excitability by activating the TrpA1 cation channel in motor neurons and induced proteomic stress in larval muscle by inhibiting ER fusion and disrupting autophagy by deletion of *atlastin* [12, 13]. We found that TrpA1 activation enhanced several phenotypes conferred by the null *atl^2^* mutation. Most notably, TrpA1 activation enhanced the *atl^2^*-dependent accumulation of muscle cytoplasmic aggregates containing ubiquitin (poly-UB aggregates). We combined the *Gal4* and *LexA* transgene expression systems to determine that *atl* loss in muscle but not neuron was responsible for this enhancement. Expressing a second excitability transgene, a dominant-negative mutation in the *Shaker (Sh)*-encoded potassium channel (*Sh^DN^*), within motor neurons likewise enhanced the poly-UB aggregate accumulation phenotype of *atl^2^*. These results suggest that increasing neuronal activity presynaptically can induce proteomic stress in the postsynaptic cell, which can be revealed in the background of certain neurodegeneration mutants. These studies thus provide one potential mechanism for mechanistic linkage between neuronal excitability and neurodegeneration.

### How might motor neuron activity induce proteomic stress in muscle?

There are several possible mechanisms by which increased neuronal excitability could induce proteomic stress in the postsynaptic cell. Two non-mutually exclusive mechanisms include increasing muscle metabolic activity and hence mitochondrial reactive oxygen species (ROS) production or increasing muscle cytoplasmic [Ca^2+^]. Increasing ROS production could accelerate formation of oxidatively damaged proteins and lead to increased production of cytoplasmic protein aggregates. Increasing cytoplasmic [Ca^2+^] has been shown to induce various toxic processes, such as activating the Calpain protease [21], and has been implicated in neuronal death occurring as a consequence of glutamate toxicity [20, 22]. Indeed, various investigators have proposed Ca^2+^ toxicity as a potential contributor to neuronal death in various neurodegenerative disorder [41, 42].

### Activity dependent proteomic stress in muscle revealed by loss of *atl*

Our data show that increasing neuronal excitability has little to no observable effect on muscle proteomic stress in an otherwise wildtype background, but significantly increases proteomic stress in the absence of *atl*. We suggest two possibilities to explain this synergistic interaction. First, increasing neuronal excitability increases protein damage in the target muscle, possibly as a consequence of increased ROS production as described above. In this view, the wildtype muscle has sufficient reserve capacity of autophagy machinery to handle this increase in protein damage without observable increases in poly-UB aggregate accumulation (Figure 8). In contrast, this reserve autophagy capacity is eliminated by *atl* loss, leading to an inability to handle the increased protein damage caused by neuronal activity. Consequently, these damaged proteins accumulate as poly-UB containing aggregates (Figure 8). Second, neuronal activity partially inhibits autophagy in the target muscle. Whereas the wildtype muscle has sufficient reserve capacity of autophagy machinery to handle this inhibition without difficulty, *atl* loss eliminates this reserve capacity and sensitizes the muscle to the further crippling of autophagy caused by increased neuronal excitability, leading to the observed enhancement of poly-UB aggregate accumulation.

**Figure 8.**
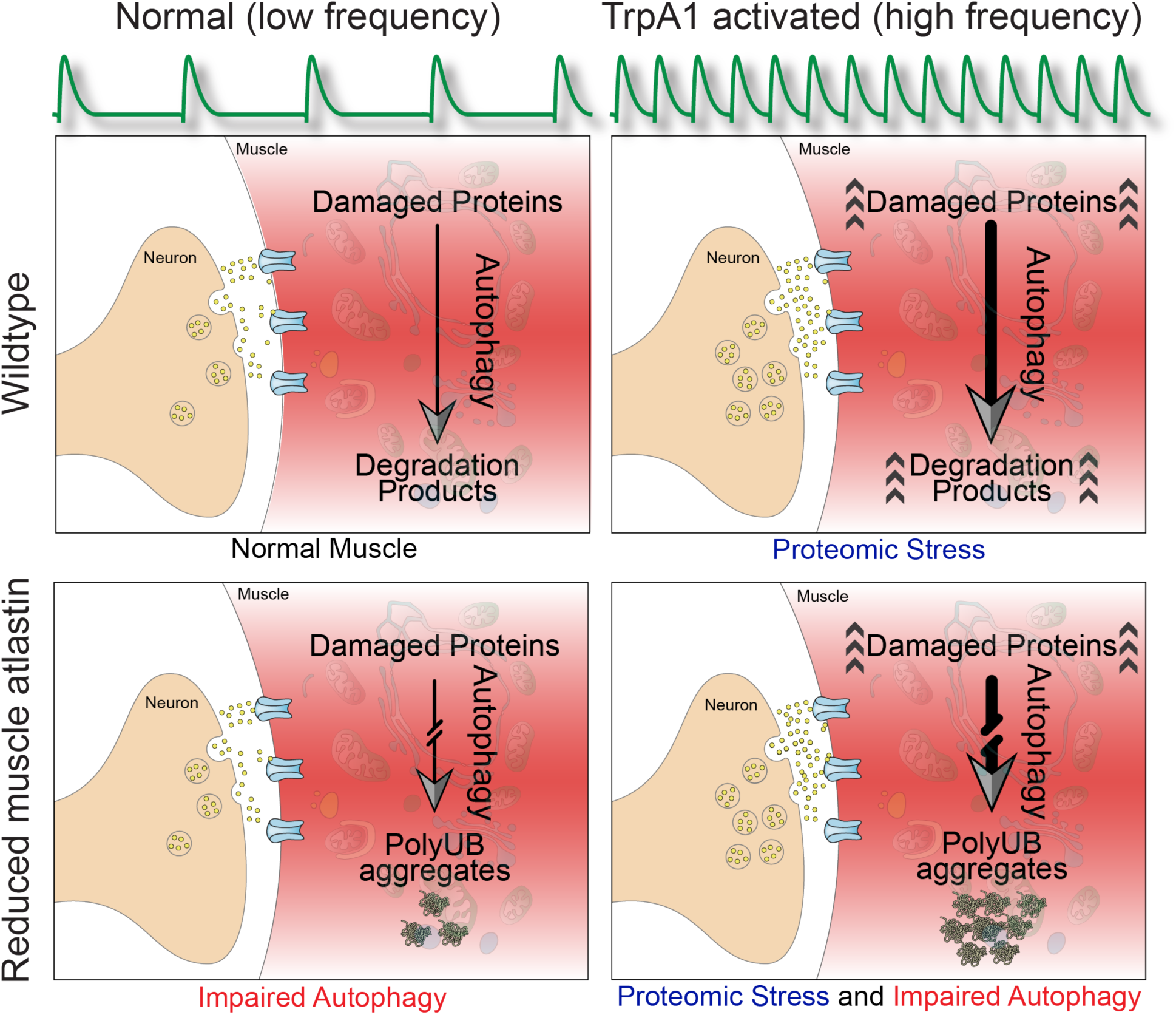
– Enhancement of the *atl^2^* poly-UB aggregate accumulation phenotype by increased neuronal excitability. At the wildtype neuromuscular junction (upper left panel), low frequency motor neuron action potentials (upper green trace) triggers fusion of synaptic vesicles (green circles) with the cell membrane, releasing neurotransmitter (l-glutamate, small green circles) into the synaptic cleft and enabling binding glutamate receptors (blue) on the surface of the muscle (pink). Damaged proteins are degraded via the autophagy pathway. In muscle lacking *atl* (lower left panel), the autophagy pathway is attenuated, leading to accumulation of protein aggregates. TrpA1 activation (right panels) increases motor neuron action potential frequency, leading to increased release of neurotransmitter and increase in protein damage (arrowheads). Whereas the *atl^+^* muscle can increase autophagy to accommodate this increased protein damage (upper right panel), the attenuated autophagy in muscle lacking *atl* (lower right panel) prevents this accommodation and leads to increased accumulation of protein aggregates.

### Relevance to neurodegenerative disorders

This work describes excitability-driven enhancement of proteomic stress specifically at the neuromuscular junction. Extrapolating these interactions to central synapses would afford relevance to other neurodegenerative disorders. In this view, neuronal activity would transition from benign to toxic as postsynaptic autophagy deficits begin and protein aggregates accumulate. Such a process could explain the well-established ability of conditions that increase excitability, such as epilepsy, to contribute to pathology of neurodegeneration. Furthermore, the observation that neurodegenerative processes can also feedback to increase neuronal excitability raises the possibility that neuronal excitability and neurodegeneration can together form a positive feedback loop, ultimately leading to exponential increases in both neuronal excitability and neurodegeneration. If so, then pharmacological interventions that depress neuronal excitability could sever this positive feedback and thus be beneficial in the treatment of a variety of neurodegenerative disorders.

## Acknowledgements

Funded by grants R01 NS102676 and R21 NS111340 from NINDS to MS and JAM. We are grateful to the Bloomington Drosophila Stock Center, Tim Mosca, Gerry Rubin and Kristin Scott for provided Drosophila stocks. This work conducted, in part, using resources of the Rice University Shared Equipment Authority.

## Figure Legends

**Figure S1.**
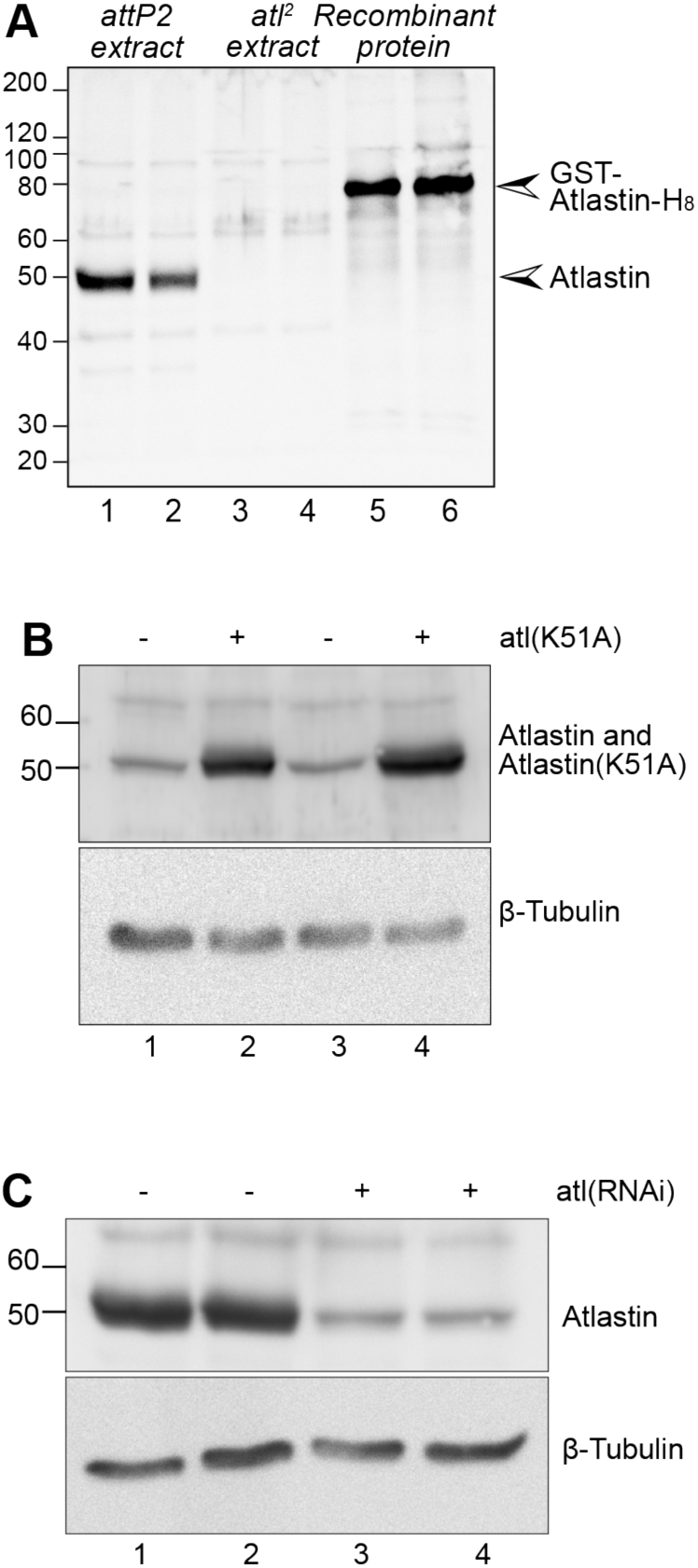
– Validation of *nSyb-lexA* driver for pan-neuronal induction of gene expression. A) Western blot with anti-Atl antibody from whole fly from wildtype (*attP2*, lanes 1 and 2), *atl^2^* (lanes 3 and 4), and recombinant GST and His-tagged Atl (lanes 5 and 6). B) Heads from flies carrying *nSyb-lexA* and either *Aop-*driven *atl^K51A^* (lanes 2 and 4) or empty vector (lanes 1 and 3). C) Heads from flies carrying *nSyb-lexA* and either empty vector (lanes 1 and 2) or *Aop-*driven *atl RNAi* (lanes 3 and 4). β-tubulin serves as loading control in B) and C).

**Figure S2.**
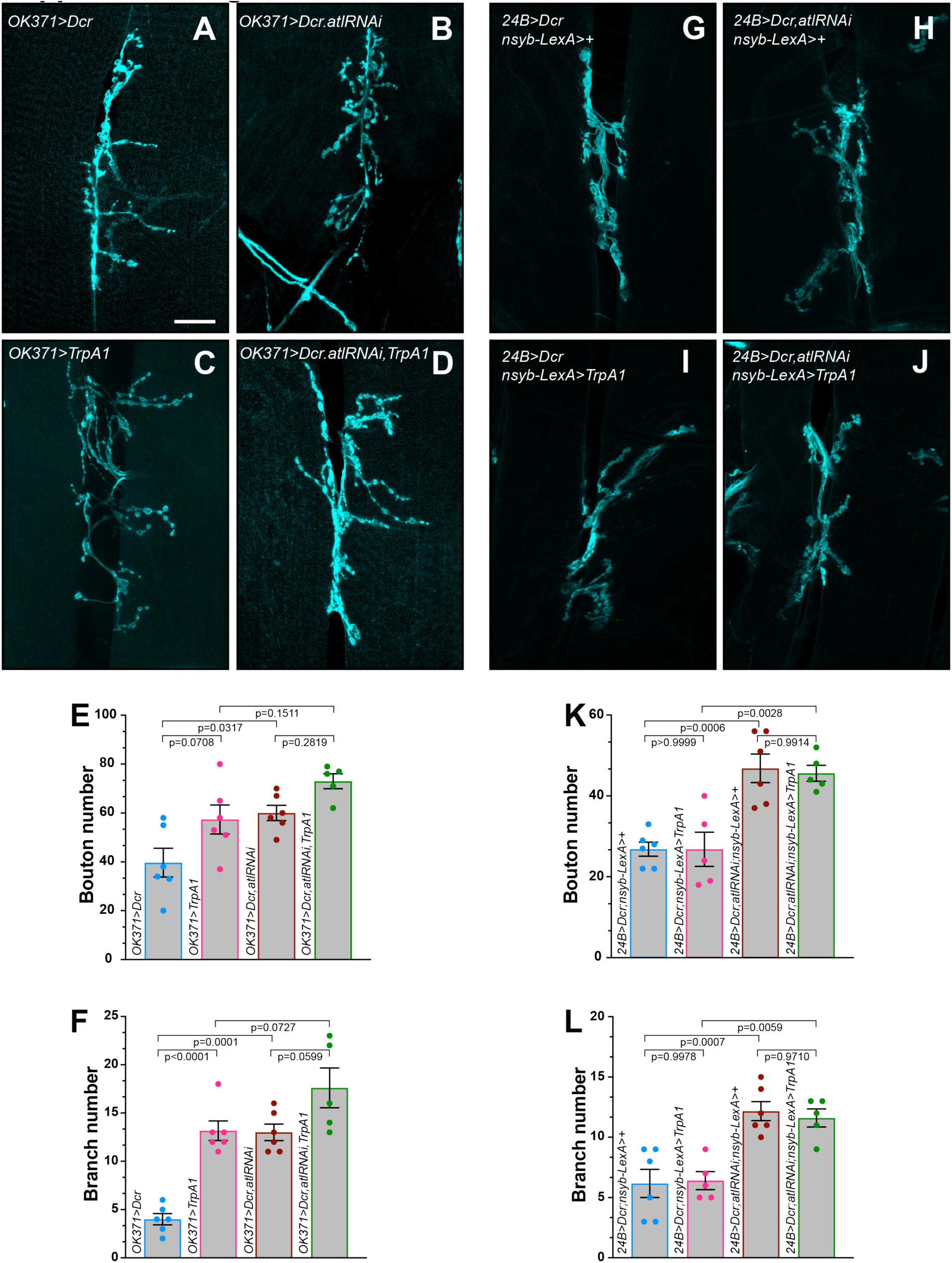
– TRPA1 activation fails to enhance the increase in synaptic bouton number and axon branching conferred by *atl* loss. Larval nerves innervating muscles 6 and 7 were labelled with Alexa fluor 647 conjugated anti-HRP and imaged on a Zeiss LSM 800 with a 40x objective. A)-D) and G)-J) show neuromuscular junctions from larvae of the indicated genotypes, shown in pseudo cyan. E), F), K), and L) Scattergram showing means +/-SEMs of bouton number (E) and axon branch number (F) for larvae of genotypes. p values were calculated by a one-way ANOVA and Tukey’s post-hoc test. n=5 or 6.

